# Raw-count embeddings improve single-cell foundation models

**DOI:** 10.64898/2026.06.29.735389

**Authors:** Sebastian Schlede, Thulasi Priyadharshini Muruganandan, Santhosh Gojjam Kantharaju, Ilmars Kisis, Maike Boecker, Mira Kim Alves Carpinteiro, Anne Schmitz, Laura Michaela Buchwald, Vignesh Sakthivelu, Gülce Sila Gülcüler Balta, Max Anstötz, Maria Adele Rueger, Roman Kurt Thomas, Filippo Beleggia

## Abstract

Single-cell transformer foundation models have grown to hundreds of millions of parameters, yet the preprocessing choices that underlie them, including gene ranking and library-size normalisation, have not been systematically benchmarked. Testing seven strategies, we find these elaborations are largely unnecessary: non-normalised, log-transformed counts give the best performance, and gene order barely matters, with even random ordering outperforming sophisticated rank-based schemes. The resulting model, Gene Intelligence, projects log1p-transformed raw counts directly onto each token embedding and jointly predicts masked tokens and counts, using no normalisation, positional encoding, or read-depth tokens. Despite this simplicity, it achieves state-of-the-art performance in the tested gene-level tasks and in doublet detection, and matches large current foundation models on cell-classification tasks while using 10-to 200-fold fewer parameters.

## Introduction

Transformer foundation models have rapidly become central to single-cell RNA sequencing (scRNA-seq) analysis. They fall into three families according to how they convert a cell’s continuous, high-dimensional counts into model input^1^. Ordering-based models rank genes within a cell by a normalised expression measure and encode the rank through gene position, simulating a natural language sentence, discarding count magnitude entirely; these include Geneformer^2,3^, tGPT^4^ and iSEEEK^5^. Value-categorisation models discretise expression into bins and embed the bin index, as in scBERT^6^ and scGPT^7^. Value-projection models add a learned projection of the expression value to each gene embedding, preserving its full resolution; these include scFoundation^8^, CellFM^1^, GeneCompass^9^ and scPrint^10^. A separate line of work replaces gene-identifier tokens with protein-language-model embeddings (UCE^11^), or modifies the attention computation to inject counts, as in Transcriptformer^12^, which omits positional encodings and adds raw counts as an attention bias under a causal mask. Notably, several models ingest raw counts but then internally normalise for library size before embedding.

## Results

We first quantified how much the rank-based preprocessing actually matters, training a transformer backbone on 96 million primary cells from the CELLxGENE Discover Census^13^ initially using six preprocessing strategies that span the natural design space for rank-based tokenisation (Fig. 1a). Three strategies sort genes by raw count and differ only in how ties are broken: randomly, by the global median transcripts-per-ten-thousand (tp10k) in expressing cells or by the gene’s rank in the global distribution. The remaining three strategies sort by fold-change against the global median tp10k, by the shift between within-cell and global rank, or shuffle genes at random. All models otherwise share an identical architecture (32,768-token vocabulary, up to 16,384-token context length, FlashAttention^14^, SwiGLU^15^ feed-forward, one cell token and six metadata tokens) and were evaluated with frozen embeddings and a logistic-regression probe on five benchmarks, including cell type classification, doublet detection and gene classification. As standard cell classification tasks have reached saturation^12^, we chose two distinct more difficult tasks. First, the identification of cell types derived independently of the expression data itself, using a Cellular Indexing of Transcriptomes and Epitopes by Sequencing (CITE-seq) dataset in PBMCs^16^ with a leave-one-donor-out scheme. Second, the classification of all cell types within the Tabula Sapiens^17^ simultaneously over the whole atlas instead of separately across tissues, using 5-fold validation split by donors. We also examined the models’ ability to classify singlets and synthetic doublets on cells randomly sampled from CELLxGENE, as well as two different gene-level benchmarks from Geneformer^3^: the classification of dosage-sensitive (haploinsufficient) transcription factors and the classification of NOTCH1-target genes.

**Figure 1:**
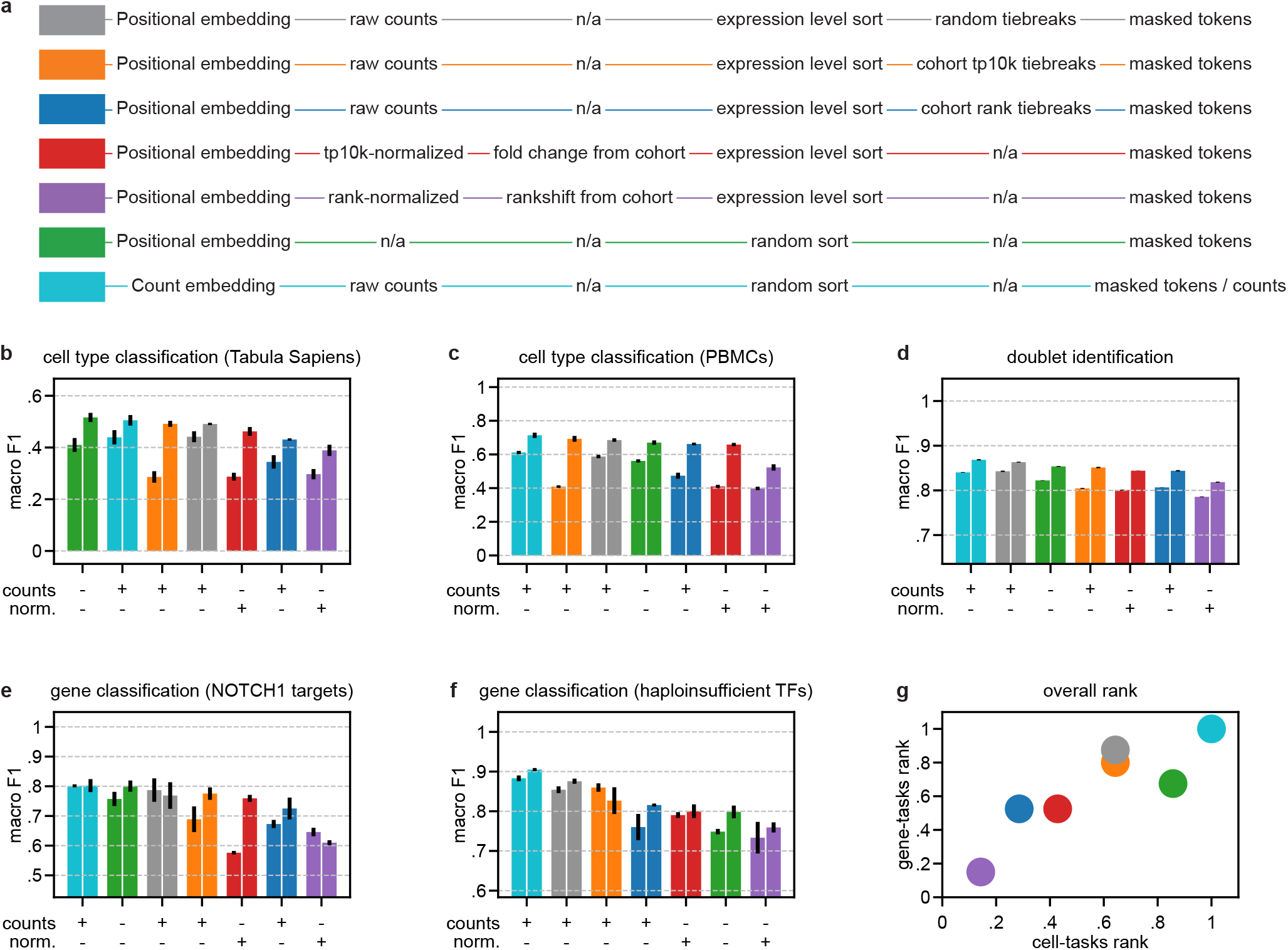
Benchmarking gene-token preprocessing strategies on five single-cell tasks. Performance of identical 12M- and 43M-parameter backbones (left and right bar, respectively) trained with each of seven tokenisation strategies and evaluated with a frozen-embedding logistic-regression probe. **a)** Schematic of the seven strategies. Each row specifies whether raw counts are used and whether they are library-size normalised, the gene-ordering rule (and tie-break), and the masking objective. The count-embedding Gene Intelligence model (teal) uses raw counts with random ordering and masks both tokens and counts, while the six positional variants encode gene order. **b)** Simultaneous cell-type classification across tissues within the Tabula Sapiens in five folds split by donor. **c)** Cell-type classification on CITE-seq PBMCs, with a leave-one-donor-out scheme. **d)** Singlet / synthetic doublet discrimination on 100,000 cells sampled from the CELLxGENE Discover Census. **e)** NOTCH1-target gene classification. **f)** Haploinsufficient/dosage-sensitive transcription-factor classification. **g)** Overall performance, shown as each strategy’s overall rank on gene-level tasks (y) versus cell-level tasks (x); the count-embedding model (teal) ranks top on both. In **b–f**, the symbols below each axis indicate whether count information without normalisation is used as the main encoder of expression level or whether normalisations are used; bars show the median across n = 5 folds for gene tasks, Tabula Sapiens classification and doublet classification and n=8 donors for PBMC classification. Error bars span the median absolute deviation from the median.

Across all five benchmarks, the count-based strategies substantially outperformed the rank-shift and fold-change variants (Fig. 1a–f). Surprisingly, gene ordering mattered little in these benchmarks, with random ordering performing best in cell-based tasks, and raw-count sorting with random tie-breaking performing best in gene-based tasks (Fig. 1g).

These observations motivated a simplified value-projection architecture without positional encodings and with a learned linear projection of log1p-transformed raw counts to each gene-token embedding. In contrast to every examined value-projection model, we applied no library-size normalisation: the projected value is the log1p of the untransformed integer count, with no depth rescaling and no auxiliary read-depth tokens. To encourage the model to use this signal, we additionally masked a random subset of counts and trained a second regression head to predict them in parallel with the standard masked-language objective. The resulting model, which we call Gene Intelligence, achieved the best score on 4/5 benchmarks and ranked first overall in both gene- and cell-based tasks (Fig. 1g), outperforming all other variants at the same parameter count.

We next compared Gene Intelligence to Geneformer, a model specialised in gene-related tasks and recently scaled to 316 million parameters^2,3^, and to Transcriptformer, a recent large model specialised in cell-level tasks and trained on multiple species (368-542 M trainable and 429-1077M total parameters)^12^ (Fig. 2a–d). On cell-level benchmarks, our 43M-parameter model surpassed Geneformer models and achieved performance equal to Transcriptformer models, despite using 10-25-fold fewer parameters (Fig. 2a,b). On both gene-level benchmarks, Gene Intelligence outperformed Geneformer and Transcriptformer by a substantial margin, establishing a new state of the art (Fig. 2c,d).

**Figure 2:**
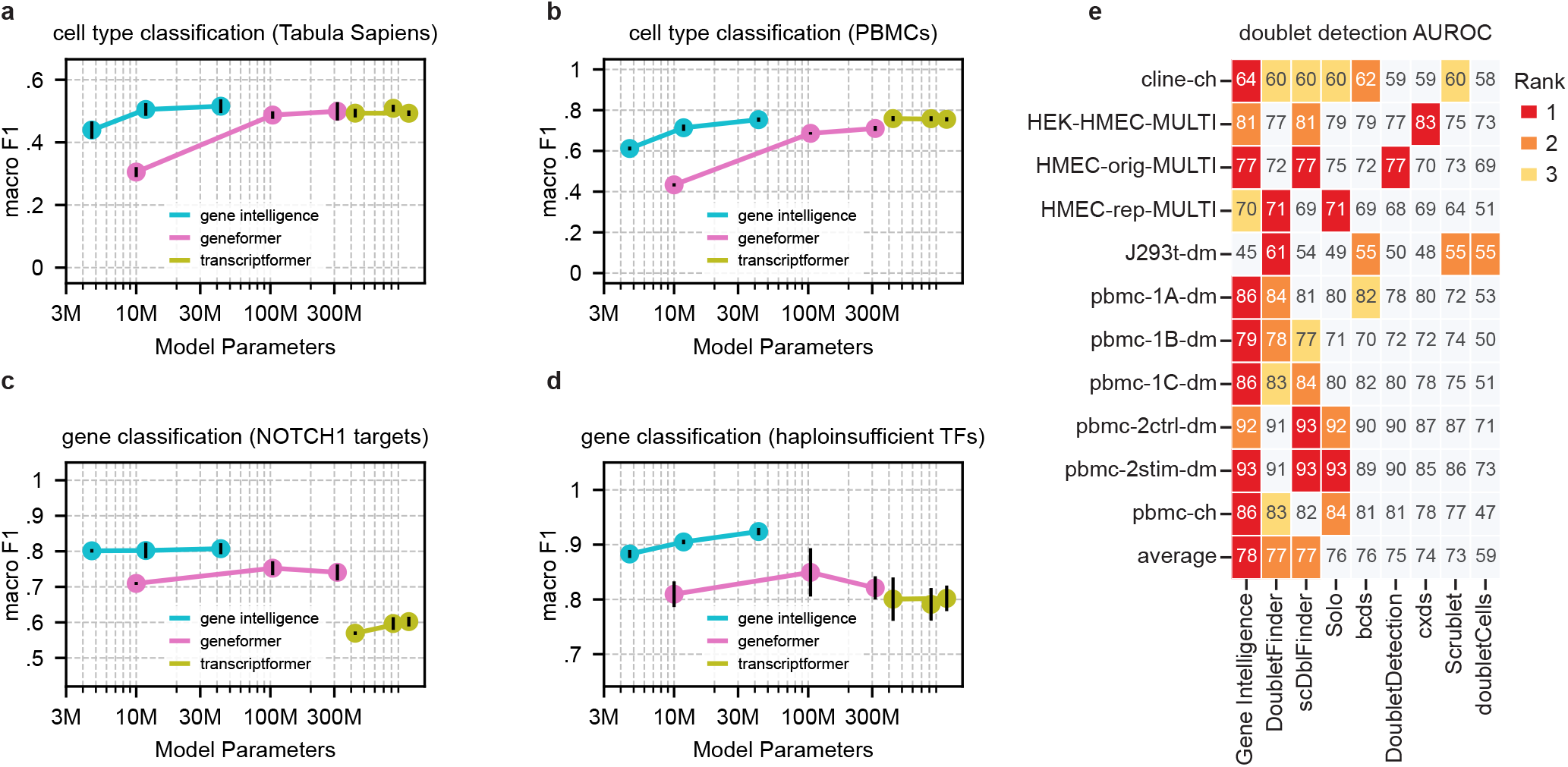
Comparison of Gene Intelligence with existing foundation models and doublet detectors across scale. **a–d)** Macro-F1 as a function of model parameter count for Gene Intelligence (teal), Geneformer (pink) and Transcriptformer (olive), evaluated with a frozen-embedding logistic-regression probe. Parameter counts include both trainable and untrainable parameters: Gene Intelligence: 4.7M / 12M / 43M; Geneformer: 10M / 104M / 316M; Transcriptformer: 430M / 824M / 1.1B. **a)** Simultaneous cell-type classification across tissues within the Tabula Sapiens in five folds split by donor. **b)** Cell-type classification on CITE-seq PBMCs, with a leave-one-donor-out scheme. **c)** NOTCH1-target gene classification and **d)** haploinsufficient TF classification. Points are the median across 5 folds for gene tasks and Tabula Sapiens classification or the median of 8 donors for PBMCs classification; vertical bars span the median absolute deviation from the median. **e)** Doublet-detection AUROC (×100) on the eleven benchmark datasets of Xi & Li, comparing Gene Intelligence to eight established methods (DoubletFinder, scDblFinder, Solo, bcds, DoubletDetection, cxds, Scrublet, doubletCells). Cell colour encodes per-dataset rank (red = best); the bottom row shows the across-dataset average, where Gene Intelligence ranks first.

To investigate the relevance of this improved performance in real-world tasks, we examined the ability of Gene Intelligence to detect doublets. Doublet detection has historically been dominated by purpose-built tools such as Scrublet^18^, DoubletFinder^19^ and scDblFinder^20^. We evaluated Gene Intelligence in this setting using the benchmark datasets compiled by Xi & Li^21^ by generating synthetic doublets within each dataset, fine-tuning the model for a single epoch on the resulting mixture, and applying the fine-tuned classifier directly without a separate probe. Across all eleven datasets that include only human cells, Gene Intelligence achieved the highest average AUROC, outperforming all eight established methods despite not having been designed for this task (Fig. 2e).

## Discussion

Three observations emerge from these experiments. First, the order in which genes are presented to the model appears unimportant, as we found that shuffling the genes randomly performed nearly as well as any principled ordering. Additionally, the most elaborate scheme we tested – reordering genes by how much their within-cell rank deviated from the cohort’s rank – was actively the worst. Second, library-size normalisation is unnecessary: projecting raw log1p counts directly, with no normalisation, positional encoding or auxiliary read-depth mechanism, leads to improved performance. Third, the resulting Gene Intelligence models are far more parameter-efficient, particularly on gene-level tasks, where our smallest 4.7-million-parameter model outperforms models more than 200 times its size.

We speculate that library-size normalisation, intended to remove sequencing-depth artefacts, also discards biologically meaningful variation in total RNA content, which reflects properties such as cell size, cycle phase and transcriptional activity. In contrast, a model given raw counts can learn the simple math needed to perform whatever depth correction a task requires while retaining this signal.

In summary, our results suggest that single-cell foundation models have accumulated preprocessing machinery – rank-based ordering, positional embeddings and library-size normalisation – that contributes little and is potentially harmful. Removing these elaborations yields a minimal count-embedding architecture that is at once simpler, smaller, and more accurate. For single-cell transcriptomics, giving a model direct access to raw counts appears more effective than feeding it the transformations the field has historically relied on.

## Methods

### Model architecture

Gene Intelligence is a pre-norm transformer encoder with SwiGLU^15^ feed-forward layers, weight-tied input/output embeddings and FlashAttention^14^ applied to variable-length packed sequences. We trained backbones at 4.7M, 12M and 43M parameter scales by jointly varying the model dimension, number of blocks, number of attention heads and feed-forward expansion. Each input sequence consists of a learned *<cell>* token, six metadata tokens (assay, suspension type, general tissue, sex, development decade, disease) and up to ~16,000 gene tokens drawn from a 32,768-entry vocabulary. Residual sub-layer projections are initialised with 1/√(2L) scaling (where L is the block index) and the prediction-head bias is initialised to the log-frequency of each gene in the training corpus.

### Gene-token preprocessing strategies

Six strategies were initially implemented in the *geneintelligence.datasets* Python module, all operating per-cell on the non-zero gene-count pairs. The random strategy shuffles all expressed genes uniformly at random. Counts / random ties sorts genes by raw count in descending order, breaking ties randomly via a pre-shuffled stable sort. Counts / global-rank ties and counts / global-median ties sort by raw count but break ties, respectively, by the gene’s rank in the global non-zero-count distribution or by the global median tp10k computed across cells expressing the gene. Rank-shift ranks expressed genes within the cell, offsets the ranks by the number of missing genes, and sorts in descending order by the difference between the cell-level rank and the global rank. Fold-change computes tp10k within the cell and sorts by tp10k divided by the global median tp10k of expressing cells. For the count-embedding variant (“Gene Intelligence”), positional encodings are disabled and each gene token additionally receives a learned Linear(log1p(count)) added to its embedding before LayerNorm and dropout, where count is the untransformed integer count with no library-size normalisation; gene order within the cell is shuffled randomly.

### Pretraining data

We trained on 96 million primary cells from the CELLxGENE Discover Census (*is_primary_data == True*; Census version 2025-11-08)^13^, restricted to datasets containing at least 10 cells and cells expressing at least 10 of the selected genes. The 32,768-token vocabulary comprises three special tokens (*<mask>, <pad>, <cell>*), the union of metadata categories across the six metadata fields, and the most abundant protein-coding and non-coding genes (ranked by the number of cells in which they are detected, with protein-coding genes detected in ≥1,000 cells given priority). Cells were tokenised on-the-fly from an LMDB index storing (gene_index, count) pairs per cell.

### Pretraining objective

Standard masked-language modelling was applied to 25 % of gene tokens; the *<cell>* token was never masked. For the count-embedding model, an additional, independent 75 % of gene-token positions had their counts masked and were predicted by a parallel scalar regression head minimising Smooth-L1 loss (beta = 1) against log1p(count). The two losses were combined with learnable Bayesian multi-task uncertainty weighting^22^.

### Pretraining optimisation

Models were trained with AdamW^23^ (β = 0.9, 0.99; ε = 1×10^−6^) using weight decay 1×10^−3^ on 2-D matrix weights and zero on embeddings, biases and LayerNorm parameters. The learning rate followed a trapezoid schedule peaking at 3×10^−4^, with 3,000 warm-up steps and 83402, 209614 and 406816 total steps, for the 2-, 4- and 8-layer models respectively. The last 16.7 % of the training steps were used for linear decay to 0.01 of the peak learning rate. Training used bfloat16 mixed precision (with float32 master weights), gradient clipping at norm 1.0, gradient accumulation to an effective batch size of 256 cells, and a token-constant length-bucketed sampler that scales the per-bucket batch size in inverse proportion to sequence length. Backbones were compiled with torch.compile and trained on a single H100 GPU.

### Benchmark datasets

Five benchmarks were used. For the two gene-level tasks, the dosage-sensitive transcription-factor and NOTCH1-target gene sets were taken from the Geneformer repository^24^. Cells for these tasks were drawn from the CELLxGENE Census: the whole Census for dosage-sensitive TF classification, and endothelial cells from the adult-heart dataset of Litviňuková et al.^25^ for NOTCH1-target classification. Cells were sampled until each labelled gene was detected in at least twenty-five cells. For the two cell-level tasks, we used cells from the Tabula Sapiens^17^, downloaded from CELLxGENE and from the Hao et al. PBMC dataset^16^. The Tabula Sapiens cells were used with the annotation provided within CELLxGENE and the PBMC dataset used the annotations provided in the source dataset. For doublet detection (Fig. 1d), 100,000 cells were sampled from the CELLxGENE Census as 50,000 singlets and 50,000 synthetic doublets, the latter formed by summing the raw counts of two randomly chosen singlets from the same pool. The separate doublet-fine-tuning benchmark (Fig. 2e) used the datasets compiled by Xi & Li^21^.

### Logistic regression evaluation protocol

For each benchmark and model, embeddings were extracted from the frozen backbone with no fine-tuning. For Gene Intelligence, cell-level embeddings concatenate the <cell> token, the six metadata tokens, and the mean-pooled gene block, all taken from the final transformer block after the final LayerNorm; metadata tokens were set to “unknown” except for assay and suspension type. Gene-level embeddings use the per-token hidden state of each gene token. In all cases a logistic-regression probe (scikit-learn, LBFGS, C = 0.01) was fit on the frozen embeddings. For the gene-level tasks, both the labelled genes and the sampled cells were partitioned into five cross-validation folds, so that each fold was evaluated on genes and cells unseen during probe fitting. For the PBMC classification task, validation used a leave-one-donor-out scheme. For the Tabula Sapiens classification task, validation used five folds cross-validation split by donors. Doublet calls (Fig. 1d) used the same probe on the 100,000-cell singlet/doublet dataset.

### Doublet detection with fine-tuning (Fig. 2e)

For each Xi & Li benchmark dataset^21^, synthetic doublets were generated by randomly merging 50 % of the cells in pairs, resulting in a final doublet content of ~33 %. Gene Intelligence was then fine-tuned end-to-end on the resulting singlet/doublet mixture for a single epoch with the standard cell-level classification head (binary cross-entropy loss; AdamW, learning rate 3×10^−4^, weight decay 1×10^−4^), and the fine-tuned classifier was applied directly to the original dataset and the published ground truth labels were used to calculate the AUROC.

### Comparison to existing methods

Geneformer^2,3^ checkpoints (10M, 104M and 316M parameters) were obtained from the official Hugging Face release^24^. Transcriptformer^12^ Sapiens (~430M parameters), Exemplar (~824M parameters) and Metazoa (~1.1B parameters) checkpoints were obtained from the official release^26^. All foundation-model comparators were evaluated with the identical logistic-regression probe pipeline applied to their respective frozen embeddings, using each model’s own preprocessing as published.

## Code and data availability

Gene Intelligence is available at https://github.com/beleggia-lab/geneintelligence. Pretrained checkpoints and configuration files will be published on publication and are available on request from the corresponding author. The CELLxGENE Discover Census^13^ is freely accessible through the cellxgene-census Python package.

